# The Golgi checkpoint: Golgi unlinking during G2 is required for correct spindle formation and cytokinesis

**DOI:** 10.1101/2023.03.05.531163

**Authors:** Fabiola Mascanzoni, Inmaculada Ayala, Roberta Iannitti, Alberto Luini, Antonino Colanzi

## Abstract

The decision to enter mitosis requires not only the control of DNA replication but also additional and crucial preparatory steps such as, for example, partial disassembly of the Golgi complex during G2. The Golgi complex is fundamental for the processing and sorting of proteins and lipids in the secretory pathway. It is organized as stacks of cisternae laterally connected by tubules to form a continuous Golgi ribbon. During G2, the Golgi ribbon is unlinked into isolated stacks in preparation for cell division. This structural reorganization is necessary for entry into mitosis, indicating that a “Golgi mitotic checkpoint” controls the correct segregation of this organelle. To understand the physiological significance of the pre-mitotic Golgi unlinking, we devised a strategy to accumulate cells in G2 with an intact Golgi ribbon and then induce entry into mitosis. Here, we show that forcing the entry of cells into mitosis with an intact Golgi ribbon causes remarkable cell division defects, including spindle multipolarity and binucleation, favoring cell transformation. We also find that the cells entering mitosis with an intact Golgi ribbon show reduced levels at the centrosome of the kinase Aurora-A, a pivotal regulator of spindle formation. Overexpression of Aurora-A rescues spindle formation, indicating that the Golgi-dependent Aurora-A recruitment has a crucial role in spindle formation. Thus, our results show that alterations of the pre-mitotic Golgi segregation have profound consequences on the fidelity of the mitotic process, representing potential risk factors for cell transformation and cancer development.

## Introduction

The G2/M transition is a critical preparatory step for the equal separation of genetic material between the daughter cells, which is crucial for cell physiology. Mitosis onset is under the surveillance of checkpoints, which are signalling pathways that arrest cell cycle progression in G2 in the presence of incomplete DNA replication or DNA damage (1). Once the checkpoints are satisfied, the irreversible commitment to mitotic entry is triggered by the activation of cyclin-dependent kinase 1 (CDK1), which induces rapid and profound modifications of cell shape, involving gradual reorganization from a flat to spherical geometry, accompanied by CE separation, nuclear envelope breakdown, chromatin condensation and spindle formation (2).

In recent years it has emerged that the control of DNA replication is not the only factor governing entry into mitosis, which also requires additional and crucial preparatory steps that occur during G2 (2, 3). Indeed, as the cells approach mitosis, the G2-restricted expression of the scaffold protein DEPDC1b induces the selective disassembly of Focal Adhesions (FAs) (4). FA are integrin-based macromolecular complexes that link the actin cytoskeleton to the extracellular matrix (5). FAs dismantling allows the retraction of the cell margin and cellular rounding by actin-based remodeling processes. Incorrect mitotic rounding alters spindle morphology and chromosome segregation (6). FA dismantling is important as its block impairs entry into mitosis (4).

An additional preparatory step is the separation of the Golgi “ribbon” into the constituent stacks. The Golgi complex (GC) has a central role in the secretory pathway (7, 8) and is generally organized as polarized stacks of cisternae connected by lateral tubules, forming a structure known as the “Golgi ribbon”(9). The current evidence suggests that an intact GC ribbon favors the efficiency of cargo processing and their polarized delivery to plasma membrane subdomains (10). Moreover, alterations of the ribbon organization are associated to modulation of signalling pathways involved in the regulation of a number of cellular processes (10–13).

We have previously shown that during G2 the GC is unlinked into stacks and that the block of this unlinking process causes a potent G2-arrest (14, 15), indicating that a mitotic “Golgi checkpoint” ensures that cells do not enter into mitosis in the presence of an intact ribbon (15, 16). The GC is the only organelle to show a structural modification in G2 as a prerequisite for entry into mitosis (9). Then, after mitosis onset, CDK1 (17, 18) triggers the disassembly of the stacks into vesicular/tubular clusters dispersed in the cytoplasm (19–21). During telophase, the Golgi clusters gradually reassemble into new Golgi ribbons in each daughter cell (22, 23).

More recently, we have also revealed the basic elements of the Golgi checkpoint. Indeed, we found that GC unlinking acts as a signal to induce Src activation at the trans-Golgi network (24), in line with the evidence that many signalling pathways are located at the GC and are modulated by its architecture to regulate high-order functions (13). Moreover, we showed that the Golgi-activated Src phosphorylates Tyr148 of the mitotic kinase Aurora-A, stimulating its recruitment at the CEs and the kinase activity (24). The latter is a novel and Golgi-specific mechanism of Aurora-A activation that is necessary for triggering mitotic entry through CDK1 activation (25).

Aurora-A activation at the CE in complex, also requiring the redistribution of a set of proteins from the FAs to the CE (26). Thus, Aurora-A acts as an integrator at the CE of stimuli from the separated GC ribbon and the dismantled FAs, suggesting that these processes are functionally coordinated to achieve the morphological rearrangements required for generating a symmetric round cell in metaphase and thus allow correct spindle formation (2).

Thus, exploiting the knowledge gained about the regulation of GC structure, we tested whether the lack of GC segregation in G2 could affect the fidelity of the division process. We asked a question different from the one addressed by Seeman’s group, which was aimed at investigating the role og stack disassembly after mitosis onset (27). The GC architecture is maintained by multiple factors, including Golgi-associated cytoskeleton (13, 28) and GC-specific structural proteins, such as the Golgi Reassembly and Stacking Proteins (GRASPs) (29) and members of the golgin family (30), which act as membrane-tethering proteins (9). The ribbon organization is characterized by a remarkable structural plasticity thanks to a dynamic equilibrium between the formation and cleavage of the inter-stack tubular connections (9, 14). Ribbon formation is driven by the Golgi “matrix” proteins GRASP65 and GRASP55 (29), which direct the tethering and fusion of newly formed tubule with an adjacent stack (9). In contrast, the cleavage of these tubules is operated by the fission-inducing protein BARS/CTBP1-S (14). The ribbon/stacks equilibrium is regulated by phosphorylation events. More in detail, during G2 a complex signaling converging on the kinases JNK2 (31), PLK1 (32) and MEK/ERK (33) induces the phosphorylation of GRASP65 and GRASP55, inhibiting their pro-ribbon role (9), resulting in unlinking (14).

Among the GRASPs, our previous studies have revealed a key role for GRASP65 because in the unphosphorylated state, in addition to promoting the formation of membrane continuity connecting the stacks, it also induces the stabilization of Golgi-associated microtubules, leading to their acetylation and clustering of Golgi stacks, which cooperate to the stabilization of the ribbon organization (34). Ribbon formation and microtubule stabilization are both inhibited by JNK-mediated phosphorylation of S274 of GRASP65 (31, 34).

Thus, to induce ribbon formation, the cells were first accumulated in G2 with an intact Golgi ribbon through JNK inhibition (31, 34). Then, the cells were forced to enter mitosis by drug-induced activation of CDK1 (35). We found that the cells showed profound alterations of fundamental mitotic processes, including the formation of multipolar spindles and binucleation. Spindle alterations were not caused by a drag force exerted on the separated CEs by an intact ribbon, as after mitosis onset the GC was disassembled also in cells treated with JNK inhibitors. The cells that entered mitosis with an intact Golgi ribbon showed reduced levels of Aurora-A at the CEs. Importantly, Aurora-A levels and correct spindle formation were recovered by Aurora-A overexpression or by inducing Golgi unlinking or disassembly, indicating that these defects were the direct consequence of the block of the GC unlinking in G2. Overall, our data revealed that the GC-originated Src-Aurora-A signaling that is triggered by ribbon unlinking in G2 is crucial for correct spindle formation in mitosis and cytokinesis and, therefore for maintenance of genome integrity. In line with this conclusion, entry into mitosis with an intact Golgi ribbon resulted in cell transformation, with important potential implications in tumor progression and maintenance of tissue homeostasis.

## RESULTS

### Setting up an approach to force entry into mitosis with an intact Golgi ribbon.

To investigate the role of GC unlinking in mitotic entry, we have developed a two-step strategy to first accumulate cells in G2 with an intact Golgi ribbon and then force them to enter mitosis. To block GC unlinking and investigate the downstream effects, we employed a well-characterized strategy involving the pharmacological inhibition of the phosphorylation of GRASP65, which induces a potent GC-dependent G2 arrest, coupled with artificial disassembly of the GC as a control to identify the direct consequences of the block of unlinking (31).

NRK cells were arrested at the beginning of the S-phase using the aphidicolin-block protocol (36) (Figure 1A). Then, aphidicolin was washed out and, during G2, the cells were treated for 2 h with dimethylsulphoxide (DMSO) as control or with SP600125 (SP), which inhibits the JNK2-mediated phosphorylation of GRASP65, resulting in the formation of a continuous Golgi ribbon and G2 arrest (24, 31). One hour after SP addition, part of the cells was also treated with MK1775 (MK), an inhibitor of Wee1, which is an inhibitor kinase of CDK1 (35). This last treatment induces the activation of CDK1 and a rapid and synchronous entry into mitosis (35). At the end of the incubation with MK, the cells were fixed and stained for immunofluorescence with an anti-GRASP65 antibody and Hoechst to label the GC and the DNA, respectively. Confocal microscopy analysis of the cells in G2 showed that the GC of control cells appeared in the usual form of distinct groups of membranes localized mainly around the nucleus, while SP inhibited GC unlinking, as previously shown (Figure 1B) (31). MK addition did not alter the GC structure of either control or SP-treated G2 cells (Figure 1B). Next, we examined if MK treatment was able to force SP-treated cells to enter mitosis. Quantification of the mitotic index showed that SP induced about 75% inhibition of entry into mitosis compared to control cells, as previously shown (31). As expected, MK addition caused a significant increase of the mitotic index in control cells. Moreover, MK also fully rescued the SP-induced G2 arrest (Figure 1C). These results indicate that MK can override the Golgi checkpoint in cells with an intact GC ribbon in G2, indicating that the assay is suitable for testing whether the Golgi status in G2 (i.e., in the form of disassembled stacks as in controls cells or intact, as in SP-treated cells) affects the subsequent mitotic processes.

**Figure 1.**
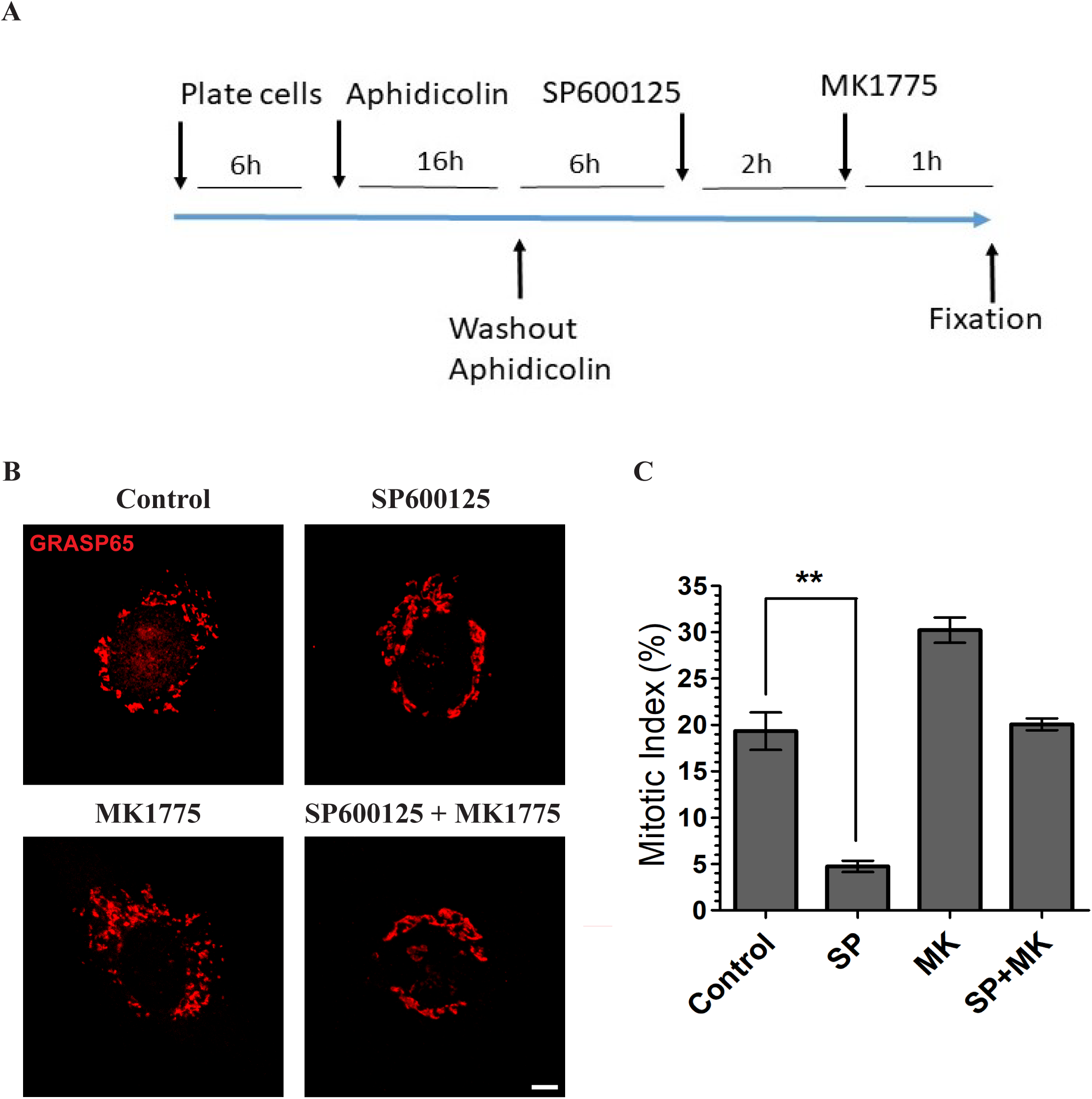
Set up of experimental strategy to induce the cells to entry into mitosis with a compact Golgi ribbon. **(A)** Schematic description of the experimental protocol. NRK cells were grown on coverslips and arrested in S-phase with overnight treatment with aphidicolin. Six hours after aphidicolin washout, the cells were treated with SP600125. Two hours later, MK1775 was added to the medium. The cells were further incubated for one hour before fixation at the mitotic peak. **(B)** Representative confocal images of the G2 Golgi complex in NRK cells treated with vehicle (Control), or the JNK inhibitor SP600125, in the presence or absence of MK1775. The Golgi complex was labeled with an anti-GRASP65 antibody. G2 cells were identified as described in (31). **(C)** Quantification of mitotic index in NRK cells treated as in (A). More than 200 cells were counted for each condition. Data are means ± s.e.m. from three independent experiments; ** indicates P<0.005, SP versus control (Control) cells (Student’s t-test). Scale bar: 5 μm.

### Golgi unlinking during G2 is required for proper spindle formation

Next, we investigated the functional consequences of the forced entry into mitosis in cells with an intact GC in G2, focusing first on the spindle structure, whose formation is completed during metaphase. The major task of the spindle is to separate sister chromatids between daughter cells. Thus, this approach offers the opportunity to clearly identify potential defects (6). To visualize the spindle aberrations, the cells were forced to enter mitosis, as indicated above. Then, after fixation, they were labeled with antibodies against α-tubulin and GRASP65 to identify the MTs and the GC, respectively, and Hoechst to visualize the DNA (Figure 2A, B). We found that 60% of the cells treated with SP and MK showed severe spindle alterations, while only a minor fraction of control cells showed alterations (Figure 2B, C), in line with the presence of transient defects normally detected also during unperturbed mitosis (37). The prevalent defect was multipolarity, with a minor fraction represented by disorganization of the spindle fibers (Supplementary Figure S1) (37). Treatment with MK or SP alone did not significantly increase the frequency of spindle defects compared to control (Figure 2C). Therefore, to test whether the observed abnormalities were a direct consequence of the inhibition of GC unlinking, NRK cells were treated as above but with the difference that 1 h before addition of SP, part of the samples were also treated with Brefeldin A (BFA) (Figure 2A) to induce artificial disruption of the GC (31, 38). BFA does not interfere with cell division and induces redistribution of Golgi enzymes into the endoplasmic reticulum (ER), while Golgi matrix proteins aggregate in the form of dispersed tubular vesicular clusters (39). Notably, the BFA-induced Golgi disassembly markedly rescued the correct bipolar spindle formation in cells treated with SP and MK (Figure 2C), indicating that a significant fraction of spindle defects was a direct consequence of an intact GC ribbon in G2.

**Figure 2.**
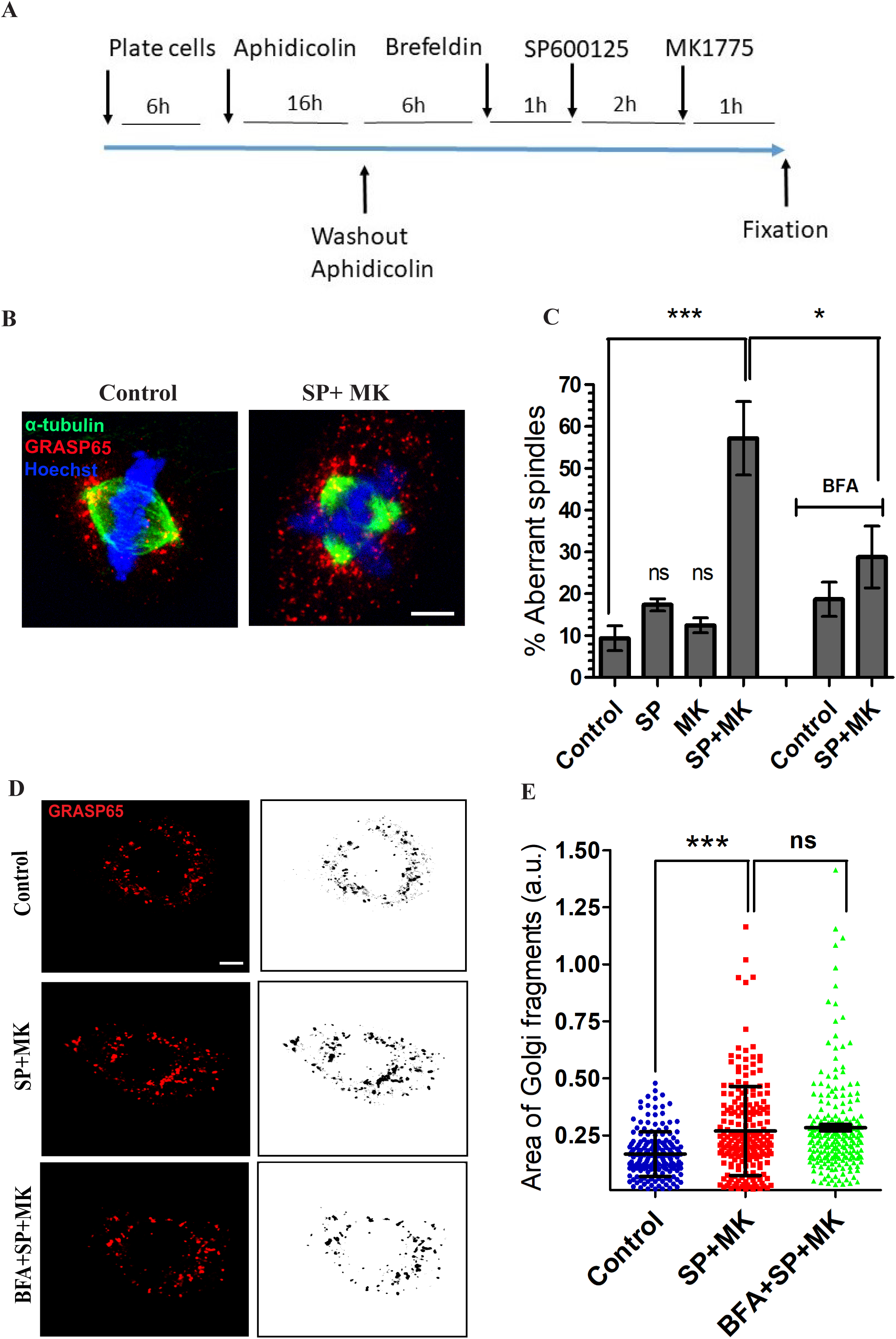
Inhibition of mitotic Golgi unlinking causes spindles defects. **(A)** Schematic description of the experimental protocol. **(B)** NRK cells were treated as indicated in (A), fixed and stained with antibodies against GRASP65 and α-tubulin to label the GC and microtubules, respectively, and with Hoechst 33342 to observe the DNA. Representative confocal images of NRK cells in metaphase treated with vehicle (Control), or the combination of SP and MK, as indicated. Control cells showed a bipolar spindle, whereas the cells treated with SP and MK presented an aberrant spindle organization (tripolar). **(C)** Quantification of the spindle defects (percentage of aberrant spindles) observed in experiments performed as described in (A). At least 200 metaphase cells were counted for each experimental condition from three independent experiments. Scale bars: 25 μm. Data are means ± s.e.m. from three independent experiments. One-way ANOVA with Tukey’s multiple comparison test. *** p < 0.001 (Control vs SP+MK); * p < 0.05 (SP+MK vs SP+MK+BFA); ns: not significant. **(D)** NRK cells were treated as indicated in (A), fixed and stained with antibodies against GRASP65, to label the GC. Representative confocal images of the GC in NRK cells in metaphase treated with vehicle (Control), or the combination of SP and MK in the absence or in the presence of BFA. For qualitative analysis of the Golgi, GRASP65 staining was processed using the ‘Invert’ function of the ImageJ software package. Scale bar, 5 μm. **(E)** Quantification of the areas of GC clusters in Control NRK cells, or in cells treated with SP and MK, in the presence or absence of BFA. Data are mean values (±s.d.) from three independent experiments. *** P < 0.0001 (Control versus cells with SP+MK) (Student’s t-test); n.s.: not significant (SP+MK versus BFA+SP+MK).

A previous study based on blocking stack disassembly through the formation of large diaminobenzidine polymers in the CG lumen, has shown that inhibition of GC stacks disassembly in mitosis induces the formation of monopolar spindles, SAC activation and metaphase arrest as a consequence of a steric hindrance of the intact Golgi stacks (27). Thus, a possible explanation of our results is that forcing entry into mitosis of G2 cells with an intact ribbon could affect spindle formation because of the persistence of large Golgi clusters, and that BFA alleviated the defects by dissolving these clusters. To examine this hypothesis, we evaluated the extent of mitotic Golgi fragmentation by measuring the size of mitotic clusters in control cells or cells treated with SP and MK in the absence or presence of BFA. In control cells, the Golgi underwent extensive fragmentation, as expected. Similarly, in cells treated with SP and MK, the Golgi was also highly fragmented, albeit to a slightly lesser extent, as shown by the about 50% increase in the average area of the GC clusters (Figure 2 D, E). The latter result is not surprising because, even if SP inhibits GC unlinking, the activation of CDK1 after mitosis onset stimulates Golgi disassembly pathways distinct from the JNK2-mediated GRASP65 phosphorylation (40). Importantly, the addition of BFA to cells treated with SP and MK did not alter the size distribution of the disassembled GC clusters compared to cells treated only with SP and MK (Figure 2 D, E), indicating that BFA does not recover spindle formation because of the reduction of the size of the clusters, suggesting that the spindle defects observed in metaphase are the direct consequence of the lack of Golgi unlinking during the G2 phase.

To test this conclusion by an independent approach, we induced constitutive and irreversible unlinking of GC into isolated stacks through the depletion of GRASP55 and examined whether this condition rescued spindle formation. TERT-RPE1 (RPE-1) cells were transfected with control non-targeting or GRASP55-specific siRNAs and synchronized using the double thymidine protocol (31). Constitutive GC unlinking is a condition that overrides the effect of SP on Golgi compaction and G2 arrest (31). After thymidine washout, the cells were incubated in the absence or presence of SP and MK, fixed at the mitotic peak and stained with antibodies against GRASP55 and α-tubulin (Figure 3A). Quantification of the results showed that in RPE-1 cells the treatment with SP or MK alone induced a minor increase of aberrant spindles. However, the combination of SP and MK had a potent additive effect in cells treated with non-targeting siRNA (about 55% of cells with defective spindles) but had a significantly lower impact in cells with the GC irreversibly unlinked by depletion of GRASP55 (GR-55 KD; about 30% of cells with defective spindles; Figure 3B). Thus, our data demonstrated that a significant fraction of the aberrant spindles observed in cells treated with SP and MK is the direct consequence of the lack of Golgi unlinking during G2.

**Figure 3.**
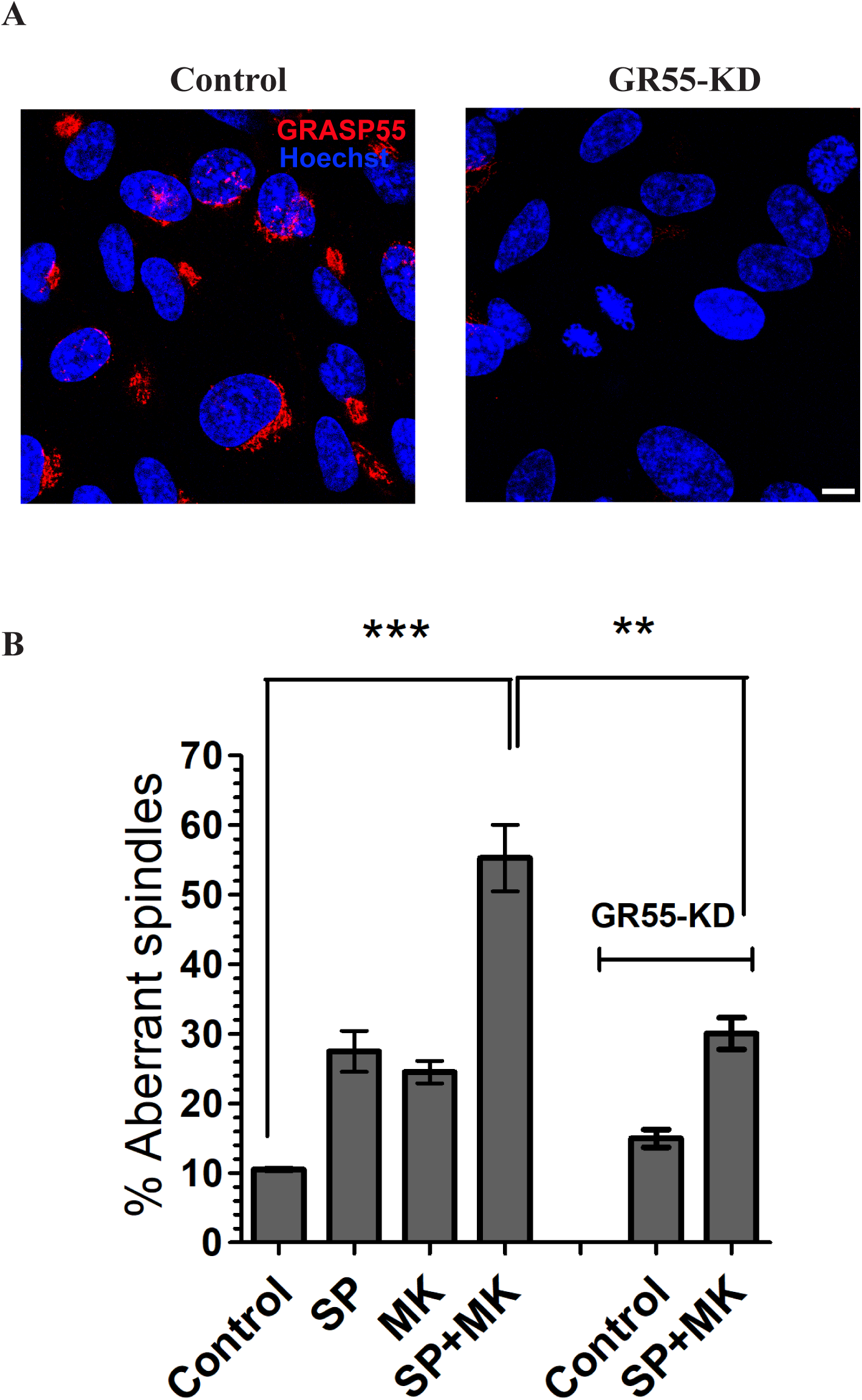
GRASP55 depletion induces a significant rescue of spindle aberration. **(A)** Representative confocal images of RPE-1 cells treated with control siRNA (non-targeting, Control; left panel) or with GRASP55-specific siRNA (GR55-KD; right panel). After fixation, the cells were labeled with an anti-GRASP55 antibody and with Hoechst 33342 to label the nuclei. Scale bars: 10 μm. **(B)** Quantification of the spindle defects (percentage of aberrant spindles) observed in experiments performed in RPE-1 cells transfected with siRNA (non-targeting, Control) or with siRNA against GRASP55, and treated as indicated. Data are means ± s.e.m. from three independent experiments. *** P < 0.001 (Control vs SP+MK); **P <0,01 (SP+MK vs GR55-KD + SP+MK). (Student’s t-test).

### Golgi unlinking controls spindle formation through Aurora-A

Next, we addressed the question of why Golgi unlinking in G2 is necessary for spindle formation. The main organizers of the spindle are the CEs, which are duplicated during S-phase (41) and then separated during G2, in concomitance with ribbon unlinking (36, 42). The separated CEs reach the definitive localization at metaphase when they are located at the opposite poles of the cells. Starting from the G2 phase, the Pericentriolar Matrix (PCM) surrounding the CEs undergoes a profound modification. The PCM is protein matrix composed of multiprotein complexes that control the polymerization and stability of three types of spindle fibers: the kinetochore MTs connecting the PMC to the kinetochores, the polar MTs that interconnect at the spindle midzone to push the spindle poles apart, and the astral MTs anchoring the PCM to the plasma membrane (41). This complex machine generates a combination of pulling and pushing forces on the chromosomes and the CEs to drive the formation of a symmetric and bioriented spindle, which is necessary for chromosome alignment at the metaphase plate (43).

The CEs are also physically connected to the GC through the polymerization of radial MTs, which form tracks along which Golgi membranes are transported to the cell center (44, 45). Therefore, it is possible that an intact Golgi ribbon could exert a drag force on the separating CEs during the G2/M transition, altering their position and thus resulting in aberrant spindle formation. To evaluate this possibility, we examined whether a bulky unseparated GC could perturb the correct separation and positioning of the CEs during G2 and prophase. To this purpose, Hela cells were synchronized for G2/M transition by double thymidine block and treated with vehicle or SP and MK, as shown in Figure 1A. The cells were then fixed and labeled with antibodies against pericentrin and GRASP65 to identify the CEs and the GC, respectively and with Hoechst to monitor the level of chromosomes condensation (Figure 4A). To measure potential perturbations of CEs separation, we assessed their “centralization” by measuring the differential distance of the two CEs from the nucleus center (Δ=Delta centralization) (Figure 4B) and CEs “separation” by measuring the distance between the two centrosomes (d=distance) (Figure 4C). Our results demonstrated that the block of Golgi unlinking did not significantly alter neither the positioning nor the separation of the CEs during both G2 and prophase (Figure 4B-C), excluding the possibility that a bulky unlinked GC could exert an important drag force on CEs repositioning. Our data are also supported by the finding that the connection between the CEs and the GC is temporarily loosened during G2 (46).

**Figure 4.**
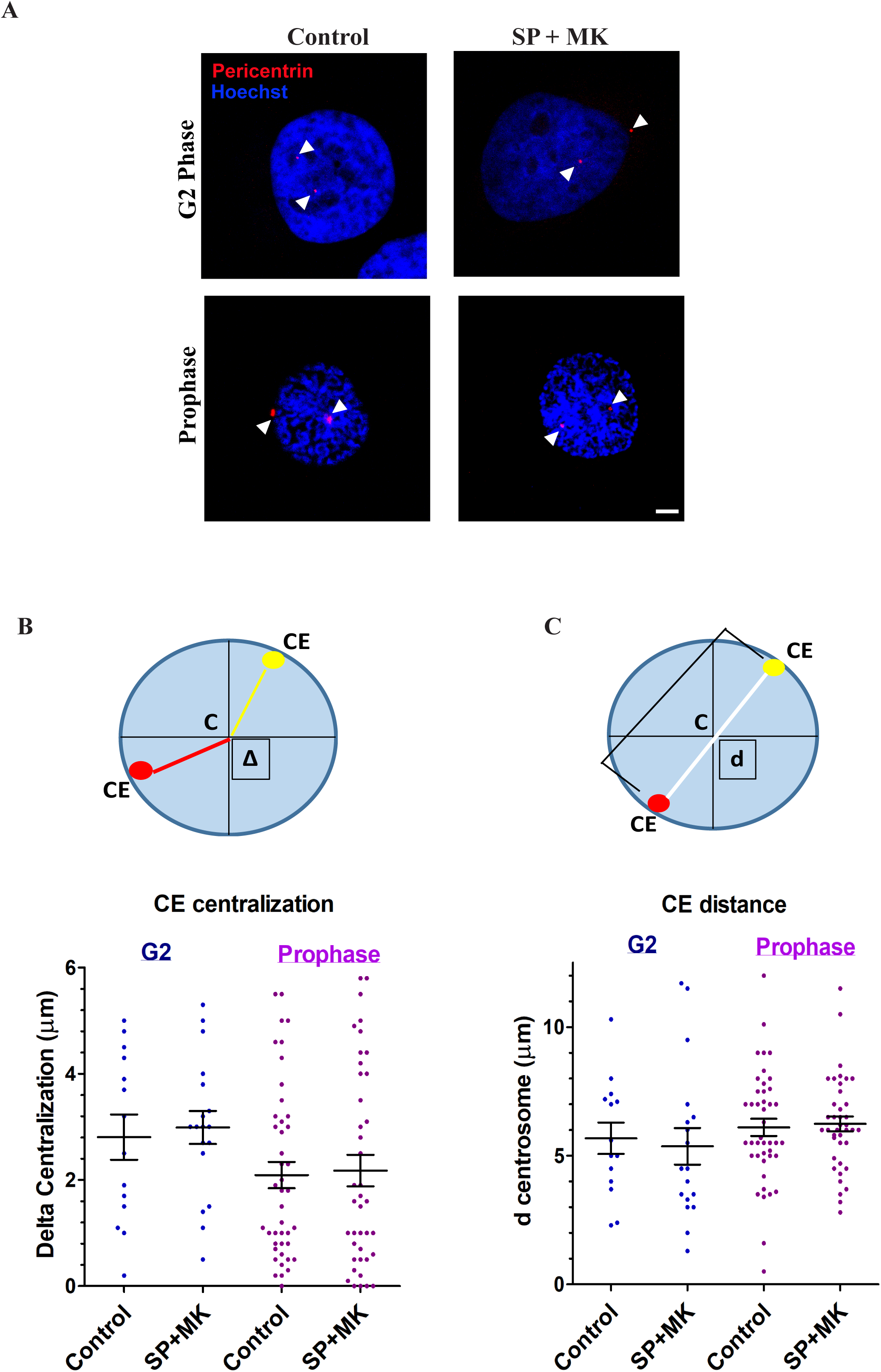
Block of Golgi unlinking during G2 does not interfere with centrosomes separation and positioning. **(A)** Hela cells were treated as described in Figure 1A and processed for immunofluorescence. Centrosomes localization was monitored in G2 and prophase by labeling the PCM with an anti-Pericentrin antibody (red) and Hoechst 33342 to label the nuclei. Scale bar, 2 μm. **(B)** Upper panel; schematic representation of the Δ parameter (Delta centralization), evaluated as the differential distance of the two centrosomes from the center of the nucleus. Lower panel: quantification of Delta centralization in cells treated or not treated with SP and MK in G2 (blue) and prophase (purple); scatter plot distribution with average value ± SEM. Student t-test showed no significant differences between Control and SP+MK-treated cells in both G2 and prophase. **(C)** Upper panel; schematic representation of the centrosomes separation, evaluated as the distance between the two centrosomes (d=distance). Lower panel: quantification of centrosomes separation in cells treated or not treated with SP and MK in G2 (blue) and prophase (purple); scatter plot distribution with average value ± SEM. Student t-test showed no significant differences between Control and SP+MK-treated cells in both G2 and prophase.

Having excluded that an intact ribbon could hamper CEs positioning in G2, we focused on the major Golgi-dependent spindle defect (i.e., multipolarity). Using confocal microscopy, we observed that this alteration is the consequence of the formation of multiple MT-nucleation foci (Figure 5A) that, in turn, are the result of the fragmentation of either CEs or PCM (43). To this end, the cells treated with control buffer and with SP with MK were fixed and labelled with antibodies against ninein (47), which is a core-CE marker that localizes primarily to the post-mitotic mother centriole and pericentrin, a PCM component (48). As shown in Figure 5B, control cells showed two pericentrin foci that colocalized with two ninein-labeled structures. Conversely, the cells treated with SP and MK showed multiple pericentrin-labeled foci but only two ninein-marked CEs. Thus, these results demonstrated that lack of ribbon unlinking in G2 causes PCM fragmentation in metaphase, resulting in the aberrant spindle formation (43).

**Figure 5.**
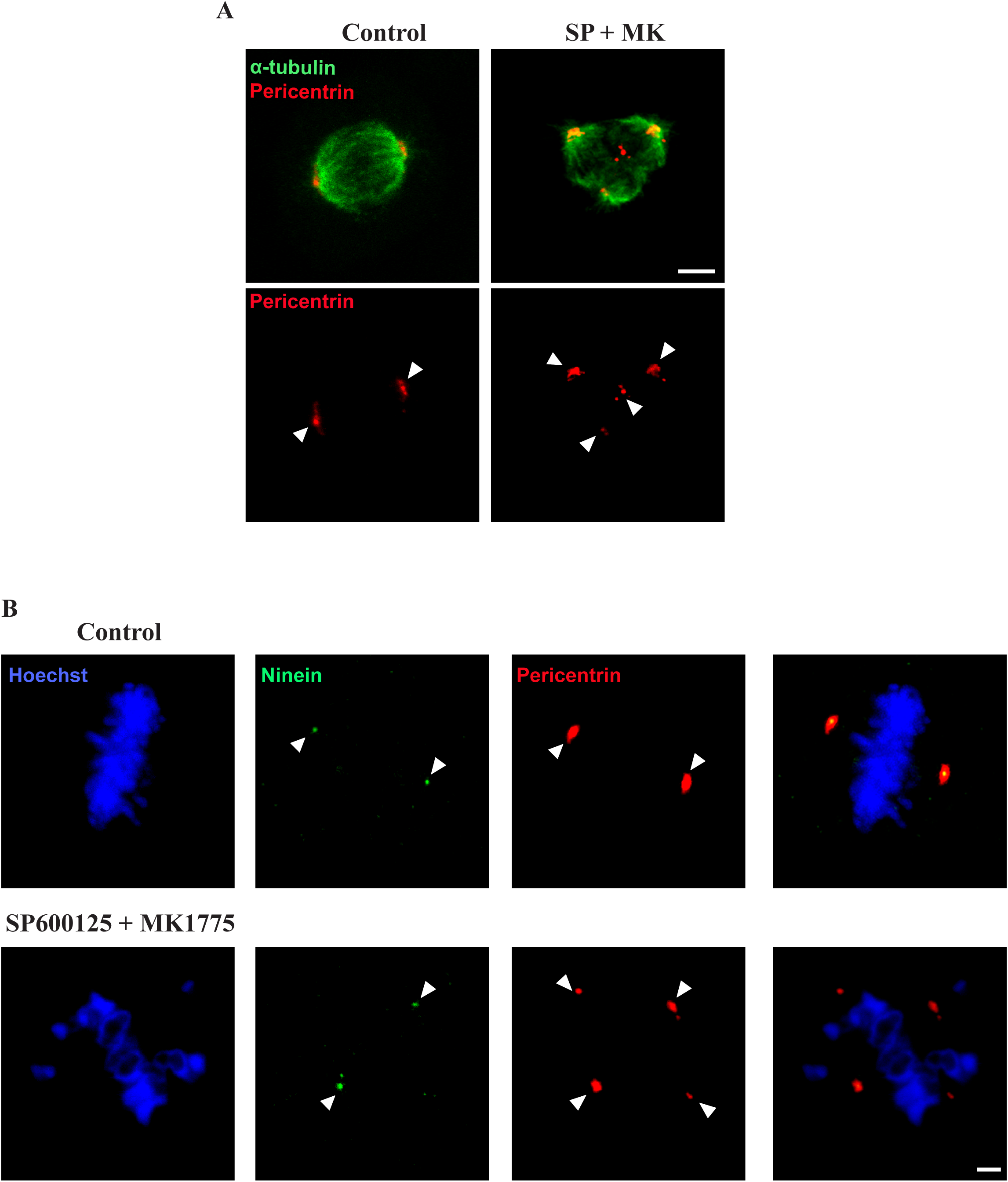
Block of Golgi unlinking during G2 causes PCM fragmentation. Hela cells were treated as described in Figure 1A and processed for immunofluorescence. **(A)** The cells were labeled with antibodies against Pericentrin and α-tubulin to label the PCM and the microtubules, respectively. The cells treated with SP and MK showed an increased number of PCM-based MT nucleation foci compared to the control cells. Scale bar, 5 μm. **(B)** The cells were labeled with antibodies against Pericentrin and Ninein to label the PMC and the centrosomes, respectively, and with Hoechst 33342 to label the DNA. Compared to the control cells, the cells treated with SP and MK showed an increased number of PCM-based MT nucleation foci, but not of centrosomes. Scale bar, 2 μm.

The fragmentation of PCM results from imbalanced forces exerted by the MT fibers on the PCMs. A major regulator of spindle fibers formation and function is the kinase Aurora-A, which during G2 becomes associated with the CEs to prepare the PCM to control MT nucleation and stabilization. The PCM comprises many scaffold proteins forming a large multimeric structure that anchors the newly nucleated MTs (49). Significantly, reduced Aurora-A activity results in altered “priming” of the PCM, leading to unbalanced MT-mediated forces during prometaphase and PCM fragmentation (49). Of note, ribbon unlinking acts as a specific signal to stimulate the activation of Src at the GC, resulting in the phosphorylation of Aurora-A at Tyr148, a residue that is distinct from the canonical Thr288 activation site. The Src-mediated phosphorylation stimulates the kinase activity and also the recruitment at the CEs (24). Thus, we tested the hypothesis that forcing the cells to enter mitosis with an intact ribbon could result in reduced Aurora-A recruitment, which is necessary for the CE-based functions of this kinase (25). The cells were synchronized for G2/M transition and treated with SP and MK in the absence or presence of BFA. The cells were fixed and stained for immunofluorescence with antibodies against pericentrin and Aurora-A (Figure 6A) to measure Aurora-A fluorescence levels at the CEs during G2 and prophase (36). Aurora-A activity can be stimulated by multiple mechanisms, including phosphorylation of sites distinct from Thr288 and binding to allosteric regulators, which operate only after its recruitment (50). Therefore, the level of recruitment at the CEs is a reliable proxy of Aurora-A activity (36, 50). Quantification of Aurora-A fluorescence intensity at the CEs during G2 and prophase showed that the treatment with SP and MK caused about a 40% reduction of its levels. Aurora-A functional activity is not directly proportional to its local concentration but requires the reaching of a specific threshold to be effective (51), indicating that the observed reduction is physiologically relevant (51). Importantly, the reduction of Aurora-A recruitment was a direct consequence of the lack of Golgi unlinking, as it was fully recovered by BFA-induced GC disassembly (Figure 6B). To examine whether the reduced level of Aurora-A at the CEs is a major cause of the spindle defects, we restored its local physiological concentration by transfecting Aurora-A-GFP. The experimental conditions were fine-tuned to induce expression levels of exogenous Aurora-A similar to the endogenous, as previously described (36). The precise control of Aurora-A expression is crucial because a high expression level can alter many mitosis-related events (25). As shown in Figure 6C and D, transfection of Aurora-A in HeLa cells synchronized for mitotic entry, induced a substantial recovery of correct spindle formation. These results indicate that the lack of Golgi unlinking in G2 causes spindle alterations because of an insufficient level of Aurora-A-activation at the CEs during G2/prophase, likely resulting in defective priming of the PCM in the control of MT polymerization or stabilization (49, 52).

**Figure 6.**
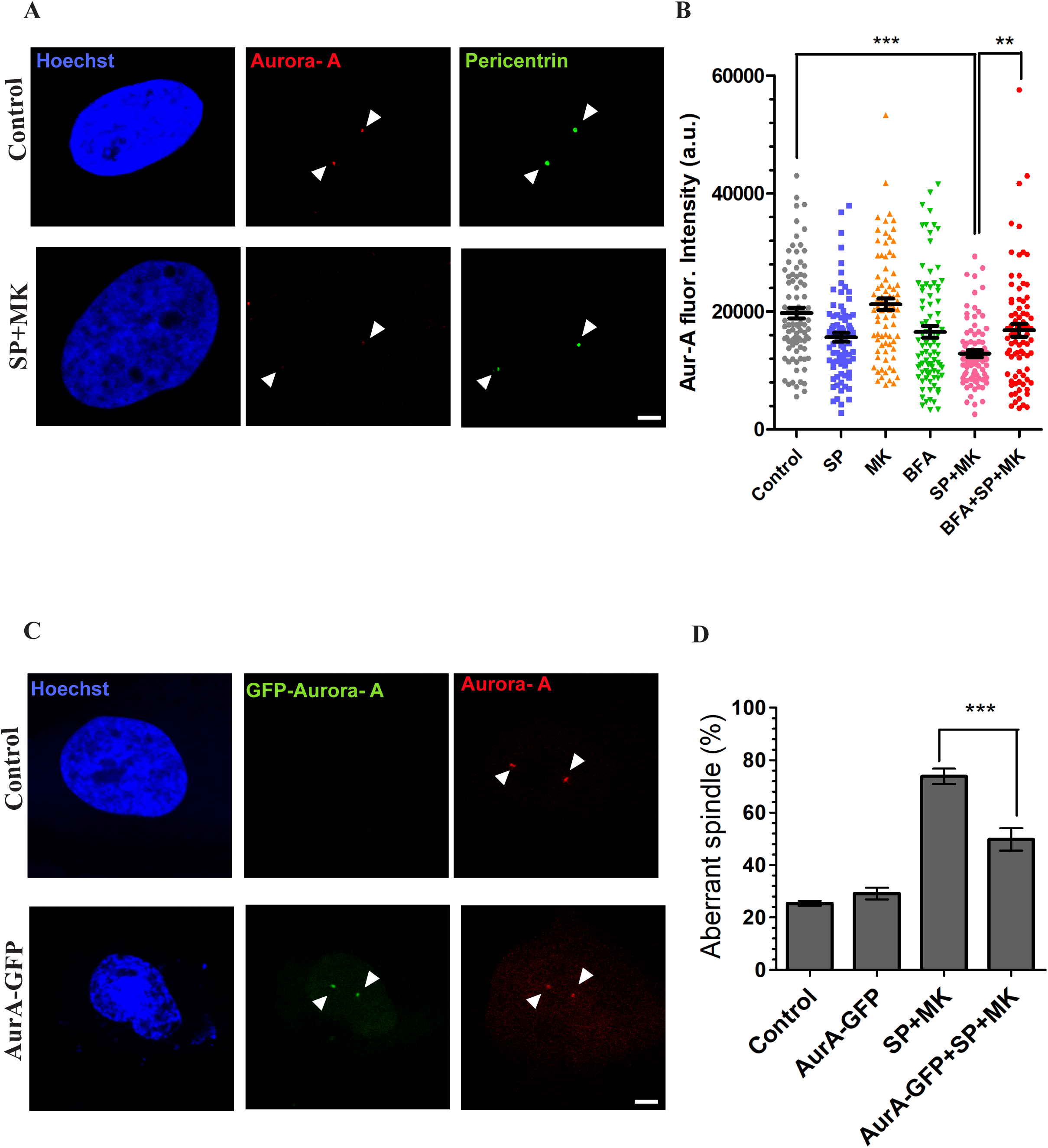
Golgi unlinking controls spindle formation through Aurora-A. **(A)** Hela cells were treated as described in Figure 1A and processed for immunofluorescence. Representative images of cells treated with vehicle or with the combination of SP and MK. The cells were fixed and processed for immunofluorescence with antibodies against Aurora-A (red), and Pericentrin (green) to label the centrosomes. Scale bar, 2 μm. **(B)** Quantification of fluorescence intensity of Aurora-A (Aur-A) in cells treated as described. All the images analyzed were acquired at maximal resolution under fixed-imaging conditions. Equal areas were used to select the centrosome regions. Quantifications are represented as scatter plots and mean values ± SEM from three independent experiments. Student’s t-tests were applied to the samples Control vs SP+MK (*** P < 0.0001) and SP+MK vs BFA+SP+MK (** P < 0,001). **(C**) Hela cells were arrested in S-phase with double-thymidine block and transfected for 24 h with the empty vector or Aurora-A-GFP after the first release from thymidine. After fixation the cells were labeled for immunofluorescence with antibodies against Aurora-A (red) and Hoechst 33342 to label the nuclei. **(D)** Quantification of the spindle defects (percentage of aberrant spindles) in cells treated with vehicle or with SP+MK and transfected with empty vector or Aurora-A-GFP. Scale bars: 25 μm. Data are means ± s.e.m. from three independent experiments. One-way ANOVA with Tukey’s multiple comparison test. *** P < 0.0001 (SP+MK vs AurA-GFP+SP+MK).

### Golgi unlinking is also required for cytokinesis

Defective spindles can result in cell death or abnormalities during mitotic exit (53). Therefore, we also examined the effects on post-mitotic events after entering into mitosis with an intact GC in G2. To this end, NRK cells were synchronized for G2/M enrichment and treated to force entry into mitosis with an intact GC, as described in Figure 2A, with the difference that the cells were fixed 16 h after the mitotic peak, as this time is sufficient to allow the duplication of most of the cells. The fixed cells were labeled for immunofluorescence with antibodies against GRASP65 and α-tubulin to detect the GC and the MTs, respectively, and Hoechst to visualize the DNA. Strikingly, confocal imaging showed that after forcing mitotic entry with an intact GC (SP+MK), more than 60% of the cells became binucleated (Figure 7 A, B). MK alone did not induce significant alterations of cell duplication. At the same time, the treatment with SP alone resulted in about 25% of binucleated cells (Figure 7B), in line with previous observations (54). Importantly, BFA-induced GC disassembly rescued cell division to levels similar to SP-treated cells, indicating that the block of Golgi unlinking is the direct cause of more than 50% of the observed binucleation events (Figure 7B).

**Figure 7.**
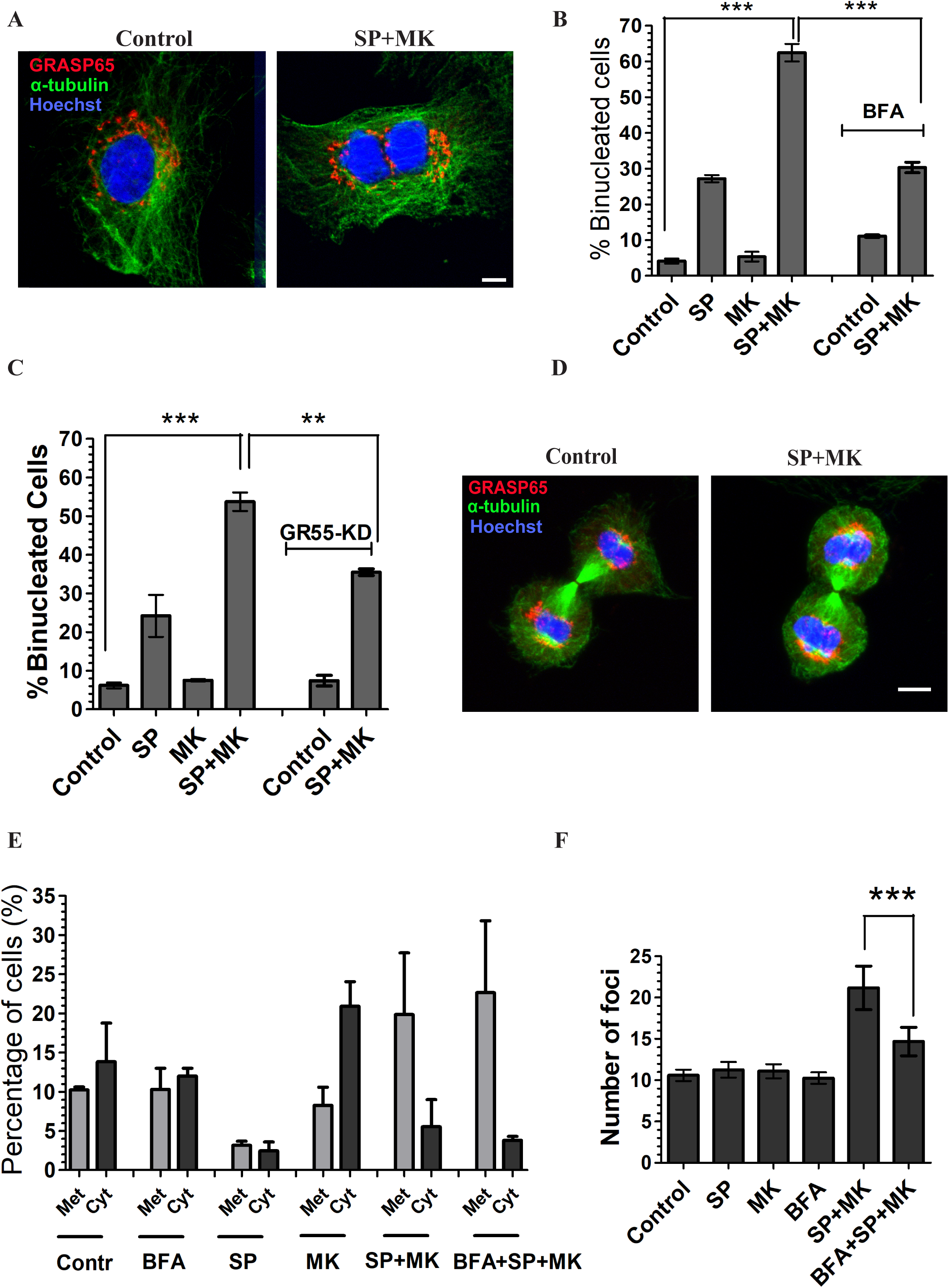
The block of Golgi unlinking causes binucleation and transformation. **(A, B)** NRK cells were treated as described in Figure 1A and fixed 24 h after aphidicolin washout. **(A)** Representative confocal images of cells treated with vehicle or the combination of SP and MK. The cells were fixed and labeled with antibodies against GRASP65 and α-tubulin to reveal the GC and microtubules, respectively, and with Hoechst to observe the nuclei. Scale bar, 10 µm. **(B)** Quantification of the fraction (%) of binucleated cells treated as described in (A). Data are means ± s.e.m. from three independent experiments. *** P < 0.0001 (Control vs SP+MK) *** P < 0,0005 (SP+MK vs BFA+SP+MK) **(C)** RPE-1 cells were transfected with control siRNA (non-targeting, Control; left panel) or with GRASP55-specific siRNA (GR55-KD; right panel) and treated as described in (Figure 3A). The fraction (%) of binucleated cells was quantified. Data are means ± s.e.m. from three independent experiments. Student’s t-test were applied: *** P < 0.0001 (Control vs SP+MK) and **P < 0,005 (SP+MK vs GR55-KD + SP+MK). **(D, E)** NRK cells were treated as described in Figure A and fixed 1,30 h after the mitotic peak. **(D)** Representative confocal images of cytokinesis (cleavage furrow formation), stained with α-tubulin to visualize the furrow, GRASP65 and Hoechst for GC and DNA, in cells treated with vehicle or with the combination of SP and MK. Scale bar, 10 µm. **(E)** Quantification of the percentage of cells in metaphase (Met) and cytokinesis (Cyt) in the experiment described in (D). Data are means ± s.e.m. from three independent experiments. **(F)** Quantification of foci formation in NIH/3T3 cells treated with control medium (CONTROL), or the combination of SP and MK in the absence or in the presence of BFA. Data are means ± s.e.m. from three independent experiments. Student’s t-test were applied: *** P = 0.0008 (SP+MK vs BFA+SP+MK).

We also tested this conclusion with an independent approach. RPE-1 cells were depleted of GRASP55 to induce irreversible Golgi unlinking (29, 55) and synchronized for entry into mitosis by the double thymidine block. The cells were forced to enter mitosis with an intact GC, as previously described, and fixed 16 h after the mitotic peak. The cells were stained with antibodies against GRASP55 and α-tubulin to label the GC and the MTs, respectively, and Hoechst to label the DNA. MK alone did not alter the binucleation index, while SP caused about 22% of binucleation. The combined treatment with SP and MK resulted in more than 55% of binucleated cells (Figure 7C). Importantly, in cells with constitutively unlinked GC (GR55-KD), the normal duplication was rescued to levels analogous to the treatment with SP alone (Figure 7C), further confirming that a significant fraction of the binucleation events is the direct result of the lack of Golgi unlinking.

On the other hand, binucleation can be the consequence of either failure of midbody cleavage or endomitosis, the latter consisting of the formation of two nuclei without cell division (56, 57). Thus, to distinguish between these two possibilities, the cells were synchronized, treated to force entry into mitosis in the absence or presence of BFA, and fixed 90 minutes after the mitotic peak. This is a suitable timing for measuring metaphase and cytokinesis events, the last-mentioned defined by the formation of the cleavage furrow. After fixation, the cells were stained with α-tubulin to visualize the cleavage furrow, GRASP65 to observe the GC and Hoechst to label the DNA (Figure 7D). The quantification of the results showed that, compared to control cells, BFA did not alter the fraction of cells in metaphase or cytokinesis (Figure 7E). Treatment with SP reduced both populations, in line with the inhibitory effect on mitotic entry. At the same time, MK increased the fraction of cells in cytokinesis, a likely consequence of the stimulation of mitotic entry. Finally, the treatment with the combination of SP and MK increased the number of cells in metaphase, indicating a delay of the metaphase/anaphase transition, with a consequential and partial reduction of cells in cytokinesis. Importantly, the addition of BFA to cells treated with SP and MK, a treatment that rescues normal cell division (see Figure 7B) did not change the numbers of cells in cytokinesis compared to cells treated with SP and MK alone (Figure 7E).

Therefore, the BFA-induced rescue of normal cell division in cells treated with SP and MK does not correlate with an increased number of cytokinesis events, indicating that endomitosis is not the mediator of the Golgi-dependent binucleation, which likely could result from failed midbody cleavage. In addition, the BFA-mediated recovery of correct cell duplication does not cause cell death, as evidenced by the counting at the microscope the number of nuclei per field of view (n/f.o.v.) 24 h after aphidicolin washout (Supplementary Figure S2), indicating that BFA does not cause an apparent recovery of cell duplication by death-mediated elimination of the failed mitotic events.

Altogether, these data indicate that the substantial increase of binucleation observed in cells treated with SP and MK could be the direct consequence of midbody cleavage failure in cells with an intact GC ribbon in G2. Binucleation can have critical pathological consequences, as the associated tetraploidization, even if transient, can lead to chromosomal instability, aneuploidy and tumorigenesis (58). In this framework, we observed that the treatment with SP and MK stimulated anchorage-independent growth in a focus formation assay, evaluated by counting the number of cell foci. SP or MK alone did not induce any effect, while forcing GC disassembly with BFA in cells treated with SP and MK strongly attenuated the anchorage-independent growth (Figure 7F). Altogether these data indicate that the alterations of the spindle and the tetraploidization caused by the lack of GC disassembly in G2 can potentially induce cell transformation.

## Discussion

In this paper, we showed that GC unlinking in G2 is necessary for the fidelity of later mitotic events, including correct bipolar spindle formation and cytokinesis (Figure 8). Indeed, forcing entry into mitosis of cells with an intact Golgi ribbon in G2 resulted in spindle multipolarity and tetraploidization. These defects were not the indirect effect of drag forces exerted by a bulky unlinked GC but the direct consequence of the lack of Golgi unlinking in G2 and of the consequently reduced activation of a Golgi-based signalling that leads to recruitment and activation of the kinase Aurora-A at the CE. Finally, our findings revealed that alterations of GC segregation in G2 could have critical pathological consequences as they can induce cell transformation.

**Figure 8.**
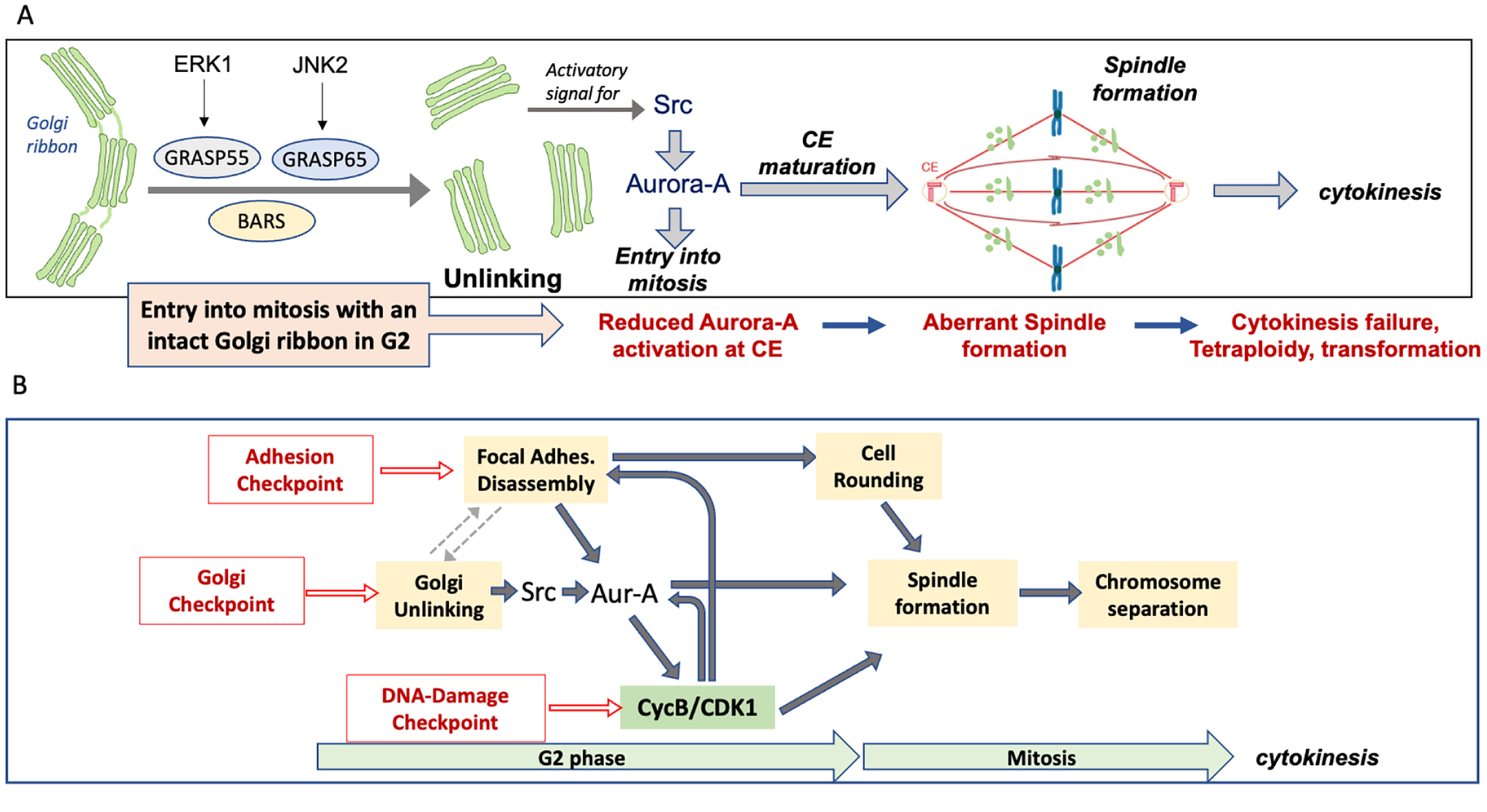
Schematic representation of Golgi unlinking and functional consequences on centrosome maturation, spindle formation and cytokinesis. **(A)** The Golgi unlinking is crucial to induce CE maturation through the Src-Aurora-A pathway activation, that leads to a correct bipolar spindle formation and cytokinesis. The block of Golgi unlinking induces aberrant spindle formation, cytokinesis failure and possible transformation. **(B)** The Golgi and the de-adhesion checkpoints coordinate Focal Adhesion disassembly and Golgi unlinking to control cell rounding and spindle formation. See text for details..

To investigate the role of Golgi unlinking in cell division, we have developed a simple strategy based on the stepwise addition of drugs and a series of controls to identify the direct consequences of the lack of GC unlinking. In brief, first, the JNK inhibitor SP is added to cells enriched in G2 to block Golgi unlinking and induce G2 arrest (31). Then, the Wee1 inhibitor MK is added to activate CDK1 and trigger a rapid entry into mitosis (35). The combined treatment with SP and MK caused severe spindle multipolarity and binucleation. Both spindle multipolarity and binucleation are the direct result of the block of Golgi unlinking as they are rescued by satisfying the Golgi checkpoint through BFA-mediated GC disassembly (31) or constitutive unlinking after GRASP55 depletion (55). Thus, our results indicate that the specific structural organization of the GC in G2 (i.e., intact ribbon or disassembled stacks) determines the correct progression of subsequent mitotic events, including spindle formation and the completion of cell abscission.

Our experimental procedure has been set up to pinpoint the role of the G2-specific GC unlinking on mitosis. Thus, it is different from the approach developed by Seemann’s group, which was based on blocking stacks disassembly after mitosis onset through the induction of large polymers formation in the CG lumen (27). Supporting the difference between the two approaches, the Golgi membranes, even if in the presence of SP, are extensively disassembled after the forced entry into mitosis with MK, thus excluding the possibility that the presence of an intact “bulky” Golgi ribbon in G2 could alter spindle formation because of a drag force exerted on CEs positioning during G2/prophase. Golgi disassembly in the presence of SP is likely caused by the activation of CDK1, which stimulates Golgi vesiculation (17). In addition, our data show that the forced entry into mitosis with an intact GC causes the formation of multipolar and disorganized spindles. Conversely, the block of stacks disassembly during mitosis causes the formation of monopolar spindles, SAC activation and metaphase arrest (27).

An important question from our data is how Golgi unlinking in G2 affects spindle formation and cytokinesis. The major components of the mitotic spindle include MT-based fibers, MT - associated proteins, and MT-organizing centers, which are often associated with the CEs, the principal spindle organizers (59). The CEs are duplicated during the G1/S transition, then, separated into two distinct CEs during G2, and reach the final position during metaphase, when they both are located at the opposite poles of the cells and are surrounded by the PCM (41). The spindle comprises three types of fibers, known as polar, astral, and kinetochore MTs, which have specialized tasks and are oriented in different directions. Many complex signals coordinate the lengthening and shortening of the various MT fibers that, together with the action of minus- and plus-directed MT motors, create a coordinated combination of forces applied to CEs and chromosomes, concurring to build a bipolar spindle with the chromosomes aligned at the spindle midzone (59). Thus, the formation of spindles involves the coordinated activity of hundreds of proteins.

Despite this complexity, our data offer a mechanistic explanation for the formation of the aberrant spindles. Indeed, investigating the role of GC unlinking in the regulation of mitotic entry, we have previously shown that ribbon unlinking acts as a signal to trigger Src activation. In turn, Src phosphorylates Tyr148 of Aurora-A, stimulating its recruitment at the CEs and kinase activity (24) through a mechanism distinct and independent of the phosphorylation of Thr288, which is the primary activation site (25). Aurora-A is essential for many distinct mitotic events, each of which is regulated by a dedicated mechanism of activation that involves the recruitment of this kinase at precise subcellular locations where it becomes active and able to phosphorylate a definite set of effectors (25, 60). In addition to stimulating entry into mitosis, Aurora-A is the primary regulator of the formation of spindle fibres (49, 52). At the G2/prophase transition Aurora-A becomes associated with the PCM, a large multimeric structure composed of scaffold proteins that organize the anchoring, polymerization and stabilization of newly nucleated MTs. The known Aurora-A substrates at the PCM include the kinase PLK1 and other scaffold proteins, including centrosomin, CDK5RAP2, CEP192, CEP215 and NEDD1, which cooperate to control MT nucleation via the γTuRC (γTubulin Ring Complex) and MT polymerization/stabilization via TACC/Ch-TOG (60, 61).

Aurora-A activity can be stimulated through multiple mechanisms, including phosphorylation of Thr288 and additional residues (e.g., Ser51, Ser98, Tyr148), binding to allosteric regulators (e.g., NEDD9, Ajuba) or by dephosphorylation of inhibitory sites (e.g. Ser342) (50, 62). Hence, due to the many different activation mechanisms, we measured Aurora-A levels at the CEs during G2 and prophase as a proxy of its activity. Moreover, Aurora-A localization is limited to the CEs during G2/prophase; thus, its levels can be unequivocally determined by immunofluorescence (36).

Notably, we found that the cells forced to enter mitosis with an intact ribbon showed reduced levels of Aurora-A at the CEs, and this phenotype was substantially recovered by artificial GC disassembly. Besides, by restoring the CEs levels of Aurora-A by transient transfection we observed a prominent, albeit not full, rescue of spindle formation. These results indicate that a major cause of spindle alterations is the insufficient level of Aurora-A-activation at the CEs due to the lack of Golgi unlinking in G2 (49, 52). However, Aurora-A over expression does not induce a full recovery of spindle formation, suggesting that additional Golgi-related events could contribute to spindle formation. Related to this issue, additional important preparatory steps during G2 are involved in cell reshaping from a flat to a spherical geometry (2). The initial steps of the rounding process begin during early prophase, but the preparatory steps occur during G2 and involve the disassembly of selected focal adhesions, which is triggered by the G2-resticted expression of the scaffold protein DEPCD1b (Figure 8) (4). Cell rounding involves the retraction of the cell margins, disassembly of stress fibers, the increase of intracellular hydrostatic pressure, and the formation of a rigid actomyosin cortex (2). The rounding process creates an optimal geometry for spindle assembly, facilitating chromosome capture by the mitotic spindle, and allows the correct orientation of the spindle, which is influenced by external cues (2). Thus, the accuracy of these architectural changes impacts mitotic progression and is essential for the proper segregation of chromosomes.

Of note, FA disassembly correlates with the relocation to the CE of scaffold proteins, such as NEDD9 and Ajuba, which protect the phosphorylation of Aurora-A in Thr288 by phosphatases (Figure 8B) (26), suggesting that Aurora-A works as an integrator of different stimuli, which are originated at the FAs and the GC, to induce PCM assembly and to activate Cdk1-CycB for mitotic entry (25) and further FA disassembly (2) (FIG 8B). Thus, the Src-Aurora-A pathway activated by ribbon unlinking could be part of a series of signaling crosstalks among the GC itself, the FA and the CEs to control spindle formation and the extensive cellular morphological reorganizations that are required to ensure proper cell rounding and thus correct spindle orientation and chromosome inheritance (Figure 8B).

Related to this model are the questions of the initial signal that triggers GC unlinking and the precise role of the unlinking process. Notably, loss of cell adhesion induces Golgi disassembly by reducing the levels of the small GTP-binding protein ARF1 at the TGN (63), indicating that FA can control GC structure and function. From the Golgi side, post Golgi traffic is specifically directed to plasma membrane hotspots juxtaposed to focal adhesions (64). Membrane traffic can transport not only cargos and membranes, but also signaling proteins (10). Thus, as a matter of future investigations, we hypothesize that FA disassembly in G2 could be the trigger of GC unlinking. Based on these considerations, and on the role of the linked GC organization in directional secretion and generation of cell asymmetry (65), we hypothesize that the unlinking process and the formation of isolated stacks could interrupt the directional delivery of membranes and signalling proteins and the generation of cell asymmetry, thus allowing the cell to reach the correct symmetry necessary for proper bipolar spindle formation. According to this hypothesis, in addition to the DNA-damage checkpoint, the entry into mitosis is controlled by the coordinated actions of the Golgi checkpoint, which is involved in the control of spindle formation, as revealed by our studies, and the adhesion checkpoint, which controls the deadhesion processes during G2. The Golgi and adhesion checkpoints could be part of a series of coordinated signalling circuits devoted to preparing the cells to the profound structural reorganizations, including the rounding process, needed to achieve the equal separation of the genetic material (2).

Overall, our work lays the foundation for future investigation of the signaling crosstalks during G2/M between FA and the GC in the control of the fidelity of mitotic events and of the segregation of the genetic material. This is physiologically relevant, as our findings show that defects in pre-mitotic Golgi segregation have significant physiological effects, as the mitotic spindle multipolarity, the consequent chromosomal instability and the cytokinesis failure cause a transient tetraploidy, which can contribute to aneuploidy (58). This latter condition promotes genomic instability, an important risk factor for cell transformation and tumorigenesis (58, 66). Supporting this possibility, entry into mitosis with an intact ribbon may favor cell transformation, as evidenced by the increased anchorage-independent formation of cell foci.

As an additional value, our findings can also open new avenues for cancer therapy, as the identification of a novel mechanism involved in controlling spindle formation can be exploited to induce mitotic catastrophe in tumor cells (67) by defining a combinatorial chemotherapeutic strategy able to target Golgi unlinking and other crucial mitotic functions, increasing drug potency and efficacy and enhancing antitumoral activity.

## Acknowledgments

The authors would like to thank the Italian Association for Cancer Research (AIRC, Milan, Italy; IG 2017 id. 20095 to AC), the POR FESR Campania SATIN for financial support and the BioImaging Facility at the Institute of Experimental Endocrinology and Oncology for support in imaging microscopy.

## FIGURE LEGENDS

**Figure Supplementary S1.**
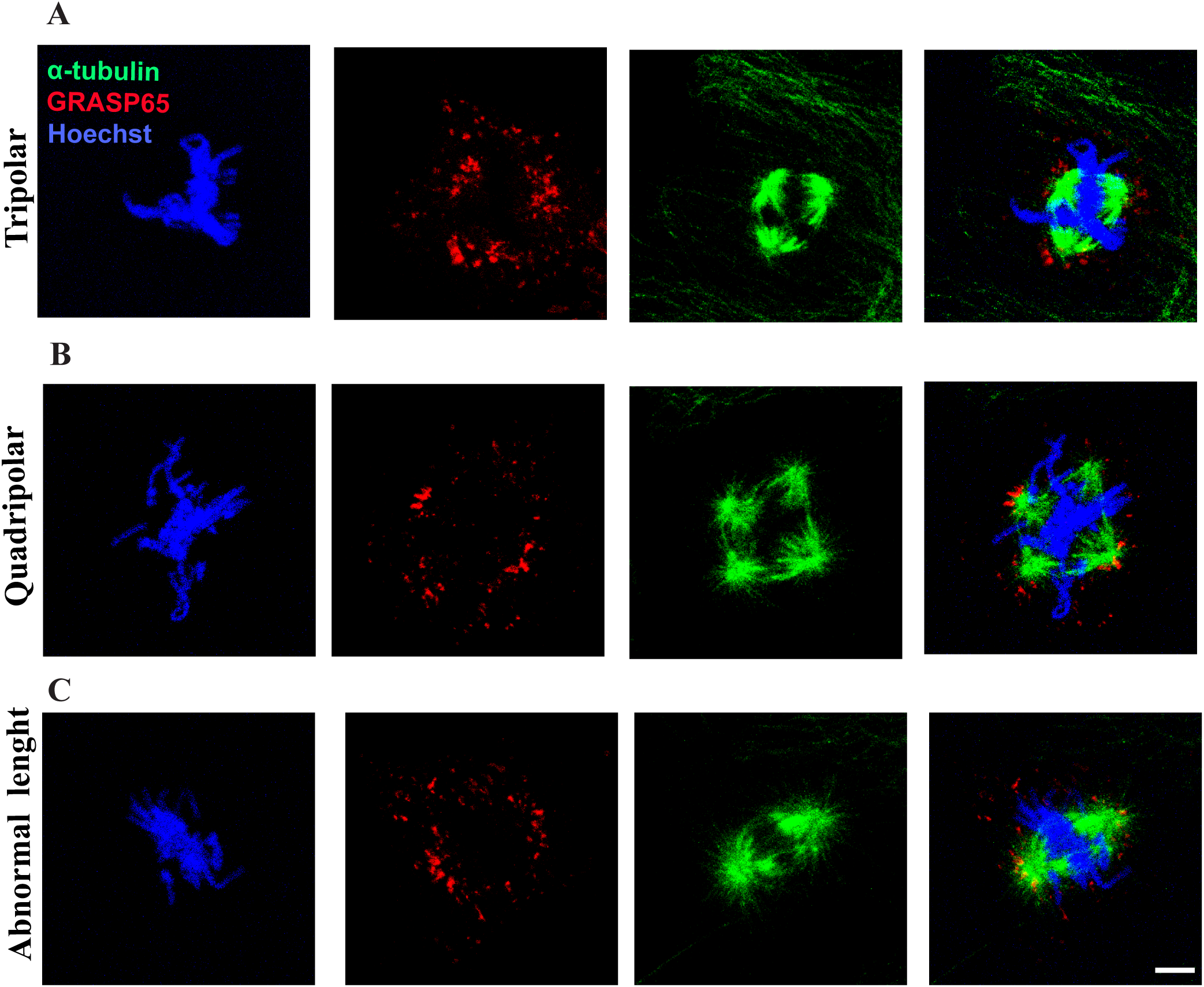
Classification of spindle abnormalities. The cells were stained with Hoechst to visualize DNA condensation, GRASP65 to visualize GC distribution and α-tubulin to visualize spindle formation. **(A)** Tripolar spindle formation, characterized by the presence of three different poles. **(B)** Quadripolar spindle formation, characterized by the presence of four different poles. **(C)** Spindle with abnormal length, characterized by the spindle length different from the normal lenght (37).

**Figure Supplementary S2.**
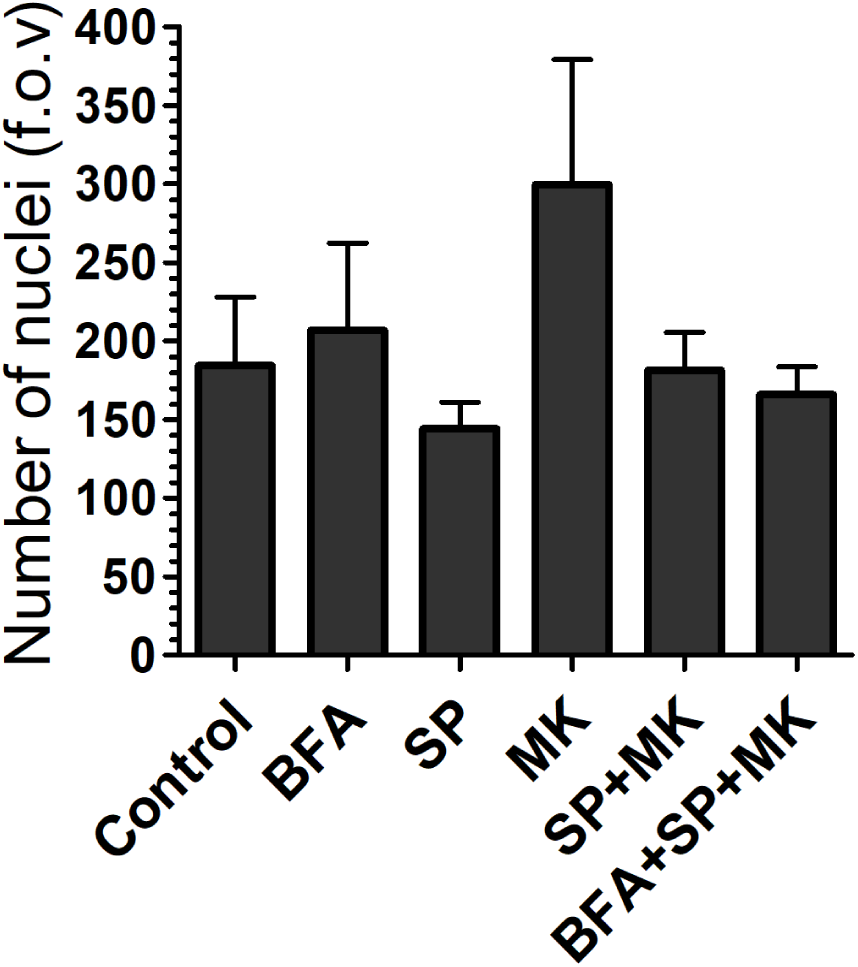
BFA does not cause cell death in cells treated with SP and MK. NRK cells were grown on coverslips and arrested in S-phase with overnight treatment with aphidicolin. The cells were fixed 24 h after aphidicolin washout. The cells were treated with combination of BFA, SP and MK, as indicated. The timing of addition of BFA, SP and MK was 5, 6 and 7 h after aphidicolin washout, respectively. The number of nuclei per fields of view (f.o.v.) was quantified. Data are means ± s.e.m. from three independent experiments. There are no significant differences between the cells treated with SP and MK and cells treated with BFA, SP and MK.

## Materials and methods

### Cell Cultures

Normal rat kidney (NRK) and HeLa cells were from the American Type Culture Collection (Manassas, Virginia) and were cultured in Dulbecco’s modified Eagle’s medium and Minimum Essential Medium, respectively (Invitrogen, Carlsbad, CA), supplemented with 10% fetal calf serum (Biochrom, Berlin, Germany), 100 μM minimal essential medium non-essential amino-acid solution, 2 mM L-glutamine, 1 U/ml penicillin, and 50 μg/ml streptomycin (all Invitrogen). hTERT-RPE1 (kindly provided by B. Franco, TIGEM) were grown in DMEM/F12 with 10% fetal bovine serum (Gibco, BRL), 2 mM l-glutamine, 1 U/mL penicillin and 50 μg/mL streptomycin (Invitrogen). NIH3T3 fibroblasts were maintained in Dulbecco’s modified Eagle’s medium (DMEM, Invitrogen, Carlsbad, CA) supplemented with 10% bovine serum (Biochrom, Berlin, Germany), 2 mM L-glutamine, 1 U/ml penicillin, and 50 μg/ml streptomycin (all Invitrogen). All the cell lines were grown under a controlled atmosphere in the presence of 5% CO_2_ at 37°C. For cell synchronization at the G2/M transition, the cells were treated as previously described (31).

### Antibodies and reagents

Aphidicolin (2.5μg/ml), Thymidine (2 mM), Brefeldin (200 ng/ml), SP600125 (25 μM) and Fibronectin (10 μg/ml) were from Sigma-Aldrich. MK1775 (0.3 μM) was from Selleckchem. DMSO was from Carlo Erba. Hoechst 33342 was from Invitrogen. Mowiol4-88 were from Sigma-Aldrich. The antibodies were from the following sources: anti-GRASP65 (1:3000) and anti-Pericentrin was from Abcam (Cambridge, UK); anti-α-tubulin (Clone DM1-A) (1:5000) were from Sigma; anti-GRASP55 was from Novus Biologicals (1:500); anti-Aurora-A was from BD Transduction Laboratories (1:200); anti-Ninein was from BioLegend (1:200). Alexa 488-, Alexa 633- and Alexa 568-conjugated secondary antibodies (1:400) were from Invitrogen.

### Plasmids, siRNAs and transfection

The GRASP55 vector was targeted using a smart pool from Dharmacon (Lafayette, California) directed against the sequences: #1 5’-GGAGUGAGCAUUCGUUUCU-3’; #2 5’-GUAAACCAGUCCCUCACUU-3’; #3 5’-GACCACACAGUGAUUAUAU-3’; #4 5’-UGUCGAGAAGUGAUUAUUA-3’. The siRNA duplexes were transfected using Lipofectamine 2000 (Invitrogen), according to the manufacturer instructions. As a NT, we used the siRNA duplex 5′-UUCUCCGAACGUGUCACGUdTdT-3′ (Sigma). The cells were further processed according to the experimental design and depletion was assessed by immunofluorescence. For transfection of Aurora-A-GFP, HeLa cells were arrested in S-phase with double-thymidine block and transfected in 24 well with 0,25 μg of vector 24 h with the TransIT-LT1 Transfection Reagent (Mirus), according to the manufacturer’s instructions. The cDNA of full-length Aurora-A fused to GFP was a kind gift from Dr Stefano Ferrari (Institute of Molecular Cancer Research, University of Zurich, Zurich, Switzerland).

### Immunofluorescence microscopy

The cells were grown on 10 μg/mL fibronectin-coated glass coverslips (Sigma-Aldrich) and treated as described above. They were then fixed with 4% paraformaldehyde (Electron Microscopy Sciences, Hatfield, Pennsylvania) for 10 min at room temperature. The blocking buffer (0.5% bovine serum albumin, 0.1% saponin, 50 mM NH_4_Cl in PBS) was then added to the cells for 20 minutes. For the labeling of α-tubulin, the cells were fixed with methanol at −20°C for 7 minutes and blocked with 1% bovine serum albumin in PBS. In any case, the samples were washed in PBS and incubated for 1 h at room temperature or overnight at 4°C with the primary antibodies in the blocking buffer. The secondary antibodies plus 2 μg/mL Hoechst were incubated for 40 min at room temperature. The coverslips were then mounted on glass microscope slides with Mowiol 4-88 (Sigma-Aldrich). The cells were fixed and stained with Hoechst 33342 for staining of the nuclei. Immunofluorescence samples were examined using a confocal laser microscope (Zeiss LSM700 confocal microscope system; Carl Zeiss, Gottingen, Germany) equipped with ×63 1.4 NA oil objective. Optical confocal sections were taken at 1 Airy unit, with a resolution of 512 × 512 pixels or 1.024 × 1024 pixels, and exported as .JPEG or .TIF files. The images were cropped with Adobe PhotoshopCS6 and composed using Adobe Illustrator CS6. High-resolution images were acquired at the Zeiss LSM880 using Airyscan detector and further processed with deconvolution algorithms of Zeiss Zen Black 2.1 software.

### Focus Formation Assay

To assess anchorage-independent growth, the parental NIH3T3 fibroblasts were cultured in six-well plate until they reached about 80% confluence. After thymidine synchronization, the cells were treated with vehicle, and SP and MK in presence or absence of BFA for 24 h. The culture medium was replaced with fresh medium every two-three days during the following 2 weeks. After staining with crystal violet, the morphological transformation was determined under a dissecting microscope.

## Quantitative analysis of GC area fragmentation

The analysis was performed using ImageJ. Images were subjected to thresholding, and the number of particles (“Golgi objects”) was calculated with the Analyze Particles function. All the images were acquired at maximal resolution, under fixed-imaging conditions. For qualitative analysis of the phenotype of the GC, the GRASP65 staining was processed with the freeware ImageJ software package using the ‘Invert’ function.

### Centrosomes Distance Parameters

Centrosomes position was determined using Imagej. Both centrosomes were tracked in G2 and prophase. Centrosome-to-centrosome distance (CE distance) was calculated by measuring the distance of centrosome foci in a single cell.

### Centrosomes Delta centralization

The differential distance from the center of nucleus of the two centrosomes was measured in G2 and prophase, using ImageJ. The center of nucleus is identified, as equidistal distance from cell edge. The distance of centrosomes from nucleus is measured, calculating the distance of each centrosome from center of nucleus.

### Fluorescence intensity of Aurora-A

The analysis was performed using Imagej. Cells were imaged with a confocal laser microscope (LSM710, Carl Zeiss; objective: 63 × 1.4 NA oil; definition: 1024 X1024 pixels). The bright centrosomal regions identified by a centrosome marker (Pericentrin) were circled, the Aurora-A fluorescence intensity in these regions and in a similarly sized background region were determined using LSM710 software (ZEN 2008 SP1), and the Aurora-A centrosomal fluorescence was calculated from these values.

## Statistical analysis

Statistical significance was determined by unpaired, two-tailed, Student’s t-test.

## Notes

### Competing Interest Statement

The authors have declared no competing interest.

